# Learning In Spike Trains: Estimating Within-Session Changes In Firing Rate Using Weighted Interpolation

**DOI:** 10.1101/041301

**Authors:** Greg Jensen, Fabian Muñoz, Vincent P. Ferrera

## Abstract

The electrophysiological study of learning is hampered by modern procedures for estimating firing rates: Such procedures usually require large datasets, and also require that included trials be functionally identical. Unless a method can track the real-time dynamics of how firing rates evolve, learning can only be examined in the past tense. We propose a quantitative procedure, called ARRIS, that can uncover trial-by-trial firing dynamics. ARRIS provides reliable estimates of firing rates based on small samples using the reversible-jump Markov chain Monte Carlo algorithm. Using weighted interpolation, ARRIS can also provide estimates that evolve over time. As a result, both real-time estimates of changing activity, and of task-dependent tuning, can be obtained during the initial stages of learning.

---

One of the essential analytic procedures in neurophysiology is the conversion of strings of observed action potential discharges (“spike trains”) into a dynamic estimate of a neuron’s firing rate. To determine whether a neuron fires more vigorously in the presence of a stimulus than in its absence, comparisons are conventionally made not between spikes within the stimulus interval, but rather between the estimated rate parameters. A wide variety of rate estimation procedures have been developed, including autocorrelation (Perkel et al., 1967), spike-triggered averaging (Pillow & Simoncelli, 2006), kernel density estimation (Shimazaki & Shinomoto, 2010), and spline smoothing (Kass et al., 2003).

Most estimation procedures produce very poor estimates given small samples, not only because they are noisy, but also because they are systematically biased. When estimation procedures are given very little evidence to work with, their implicit assumptions can dominate the estimate. The most common bias is dramatic attenuation around abrupt transitions in firing, such that the true rate consistently falls outside the estimate’s confidence interval. These limitations are keenly understood by analysts, who have historically performed their analyses on large sets of data to avoid these problems.

A consequence of this assumption is that traditional spike train analyses are ineffective at uncovering signals that change over the course of the recording session. This effectively makes the study of the neural codes associated with learning itself impossible. Instead, researchers typically divide spike rasters into pre-and post-learning epochs, based on behavioral performance criteria (e.g. Paz et al., 2003; Satoh et al., 2003). Thus, the vast majority of studies purporting to study the electrophysiology of learning, as well as simulations of neural networks (Amit & Brunel, 1997), are only able to reveal the consequences of learning, not the real-time processes that give rise to learning.

Despite this, there is persuasive evidence that firing rates can change abruptly during learning. Gale et al. (2014) report spike rasters with trial number as the explicit vertical axis, and show pronounced changes in firing near the time of a qualitative shift in performance. Morrison et al. (2011) also report changes in activity immediately following an abrupt task change.

In this paper, we propose a new method called *Adaptive Rate Regression Involving Steps* (ARRIS), which offers several advantages over existing methods. Chief among these is that ARRIS performs well when estimating firing rate using very few trials. Because ARRIS does not require large numbers of trials to obtain a reasonable estimate, it offers the possibility of examining how firing changes over short intervals.

Full implementation of ARRIS requires several techniques. The following list outlines the steps needed to obtain instantaneous estimates of firing at each point in a session, rather than estimates marginalized across entire sessions that are presently reported.

- A brief survey of current methods demonstrates that even very sophisticated methods perform poorly on small samples. Of these, the most promising candidate for modification is BARS (DiMatteo et al., 2001).
- BARS is effective thanks to a procedure called ‘reversible jump Markov chain Monte Carlo’ (RJMCMC). ARRIS makes unattenuated estimates in small samples by using step functions instead of the regression splines used by BARS.
- When trials share features in common, they may be combined in a weighted fashion. Weighted interpolation permits estimation of activity at specific times within a session, rather than across the session as a whole. This allows learning to be identified in real time, rather than being limited to a pre-learning/post-learning dichotomy.
- Using a temporal weight function introduces the problem of specifying an appropriate bandwidth over which to smooth the estimates. The method of generalized cross-validation (GCV) provides a metric for evaluating which bandwidths would be optimal at each time point.
- Finding the optimal GCV score must be done numerically, and this interacts poorly with RJMCMC’s reliance on stochastically approximate posterior distributions. A kernel density approximation of ARRIS can instead be used to identify which bandwidths to give the weight function.
- When ARRIS is run using GCV-optimized bandwidths for the weight function, it yields a reliable estimate for the firing rate at each time point in a session, thus revealing real-time changes in firing dynamics.

## Limitations In Existing Methods

One of the standard methods for evaluating the efficacy of density-or rate-estimation procedures is by evaluating the *mean integrated squared error* (MISE) (Silverman, 1986) of a given fit against its generating function. This entails averaging over all discrepancies between the true function and the estimate, which is straightforward when using simulated data. Figure 1 showcases how well four rate estimation procedures estimate a simple step function, given three different samples. Spikes in these simulations were generated by an inhomogeneous Poisson process, firing at 25Hz between 1s and 3s and firing at 5Hz otherwise. The samples differed in size (consisting of 5, 25, or 125 trials), and while better estimates are always expected as the sample size increases, it is also important to consider whether an algorithm can produce reliable small-sample estimates. To facilitate this comparison, the absolute error of the estimate is also reported, as are the mean integrated error and MISE.

**Figure 1:**
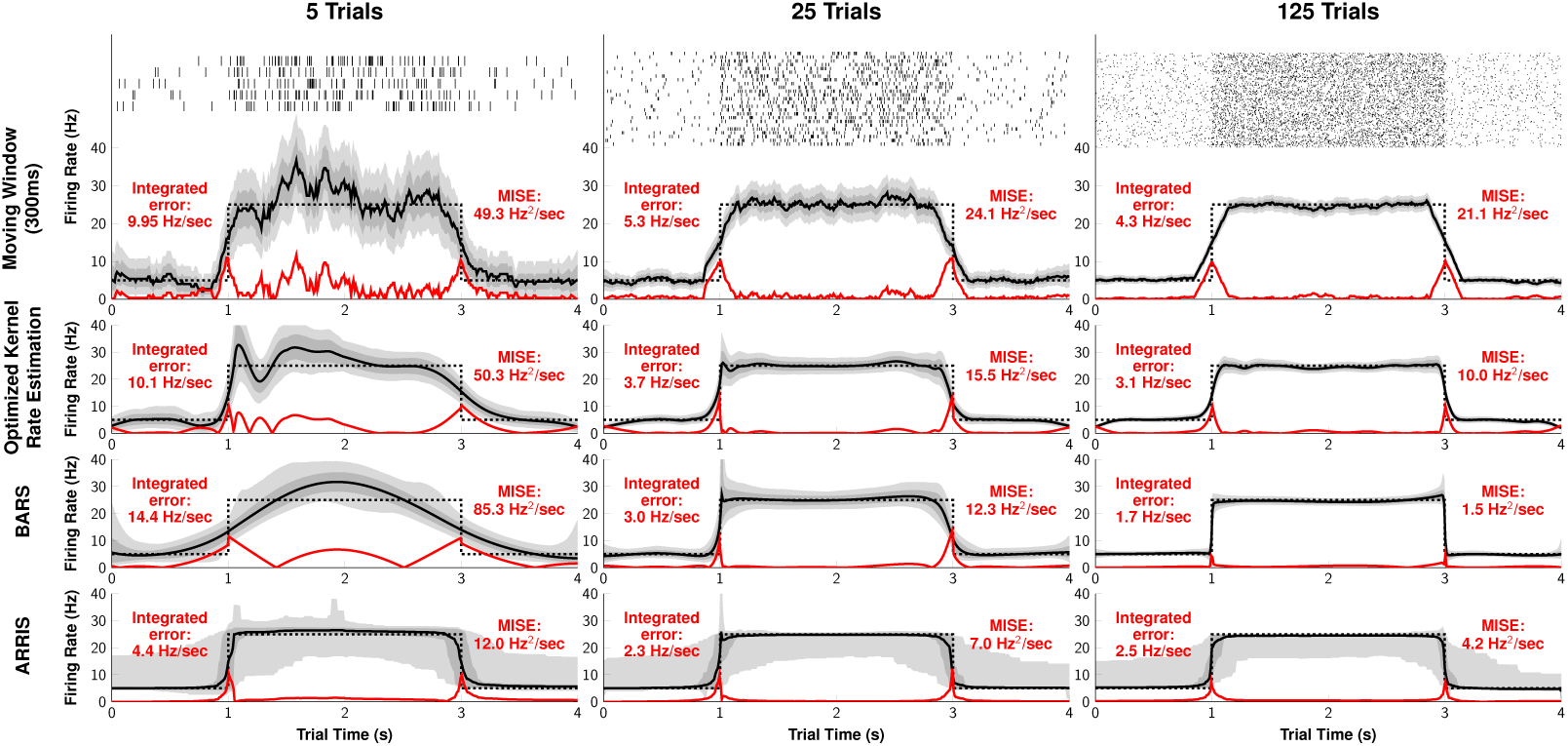
Firing rate estimates based on samples of varying size. Spikes were generated by an inhomogeneous Poisson process (dotted line). Estimate are plotted as black solid lines and shaded regions represent the 80% and 99% credible intervals of the estimates. The red line depicts the absolute error (i.e. the distance between the true rate and the estimate) at each time point. The spike rasters used for all estimates are depicted at the top of each column. **Row 1**. Rectangular smoothing function over a 300ms moving window. **Row 2**. Kernel rate estimate with an optimized variable bandwidth, using the method of Shimazaki & Shinomoto (2010). **Row 3**. BARS estimate, using the methods of DiMatteo et al. (2001). **Row 4**. ARRIS estimate, using methods described below.

Row 1 shows smoothing using a 300ms moving window, which is noisy and unreliable. Although none of its errors endure for very long, they nevertheless contribute a constant factor. Importantly, the two abrupt changes in firing (at 1s and 3s) produce unavoidable attenuation even when hundreds of trials are included. Thus, using an arbitrary fixed moving window will always produce an incorrect estimate, insofar as any rapid changes will necessarily be attenuated. This attenuation can be reduced by narrowing the width of the moving window, but this also increases the noisiness of the estimate.

A moving window estimate can also be interpreted as a kernel rate estimator, where the width of the window corresponds to a “bandwidth” parameter. Kernel rate estimators replace each observation with a “kernel,” and an estimate of the rate is achieved by summing the kernel densities. Our choice of 300ms was arbitrary, and another choice could have been made to optimally minimize the MISE. However, no single bandwidth is simultaneously optimal for both the abrupt transition and the intervening stable periods.

Row 2 demonstrates a different kernel rate estimator described by Shimazaki & Shinomoto (2010), using a Gaussian kernel. Their approach allows the bandwidth of the estimator to change over time, letting it fit both abrupt transitions and broad steady rates. This results in a systematically better fit (and lower MISE) when many trials are included. However, even when using a kernel smoother with an optimized variable bandwidth (row 2), small attenuation effects occur around transition points. Thus, although kernel smoothers are *asymptotically optimal*, they will display attenuation effects that scale as a function of sample size. For even moderately sized samples consisting of dozens of trials, this attenuation is large enough to predict anticipatory increases in firing rates (e.g. prior to a stimulus onset) that are entirely artifactual.

Row 3 fits the data according to a powerful method for spline regression, entitled *Bayesian adaptive regression splines* (or “BARS”) (DiMatteo et al., 2001; Kass et al., 2003; Wallstrom et al., 2008). The splines used by BARS are governed by “knots” whose positions control the curvature of the function. To achieve a more complex curve, more knots are required. Crucially, BARS not only provides an estimate of the firing rate, but also provides a full posterior sampling distribution for the rate estimate at each time point. That is, BARS effectively tests a representative sample of all the different splines that *could* generate the data, and reports the average of this set of estimates. This permits, among other things, a rate estimate that includes credible intervals (Chen & Shao, 1999) without relying on parametric assumptions.

BARS fares substantially better than kernel-based methods with moderate-to-large samples. Abrupt transitions remain somewhat problematic, insofar as spline regression cannot model discontinuities directly. Nevertheless, for large samples, BARS not only correctly estimates the function, but does so with very high confidence. The same cannot be said for its estimate based on five trials, which shows *more* attenuation than the kernel estimates do. This, again, is a result of the tradeoff faced in any model selection paradigm: Since BARS uses splines, and since splines assume gradual changes in rate until overwhelming evidence suggests otherwise, attenuation is an inevitable consequence when samples are small.

Dramatic attenuation of this sort is rarely reported because it is understood that unbiased estimates require that the data be pooled across many trials. Spike rasters that depict activity of single neurons typically do so by stacking multiple trials vertically, often doing so with no label on the vertical axis (e.g. Hahnloser et al., 2002; Gelbard-Sagiv et al., 2008; Jacob et al., 2013). In other words, rate estimation procedures ordinarily treat trials as *exchangeable*, having no intrinsic ordering. The estimated firing rate is then a kind of marginalized average over the full raster.

Pushing this example to its lower limit, Figure 2 gives five examples of each algorithm’s performance given spikes during a single trial. The fixed moving window (row 1) displays continuous poor performance, whereas an kernel rate estimation (row 2) and BARS (row 3) both take the form of slowly-undulating curves that heavily attenuate the source function.

**Figure 2:**
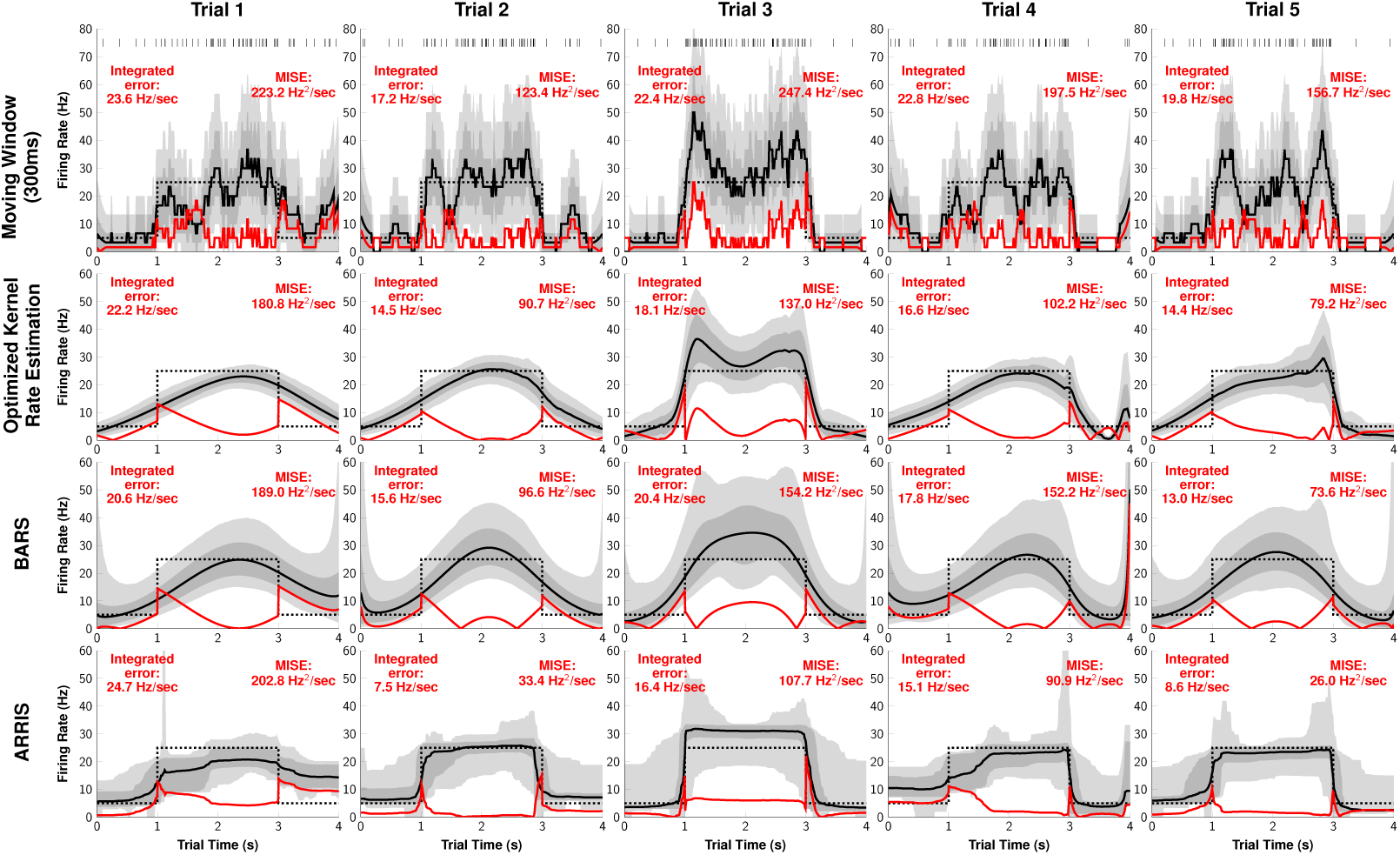
Firing rate estimates based on single trials. Spikes were generated by a Poisson process (5Hz, 25Hz, 5Hz) plotted as a dotted line. Each algorithm’s estimate of the firing rate is plotted as a solid line. Shaded regions represent the 80% and 99% credible intervals of the estimates in all cases. These were obtained from bootstrap resampling of the original data (for rows 1 and 2) or directly from the simulated posterior distributions of model estimates (rows 3 and 4). The red line depicts the absolute error. **Row 1**. Rectangular smoothing function over a 300ms moving window. Also included are the raster of spikes used for all estimates in each column. **Row 2**. Kernel rate estimate with an optimized variable bandwidth, using the method of Shimazaki & Shinomoto (2010). **Row 3**. BARS estimate, using the methods of DiMatteo et al. (2001). **Row 4**. ARRIS estimate, using methods described below.

Row 4 in Figures 1 and 2 are estimates obtained with our novel ARRIS procedure, which is closely related to BARS. ARRIS has the lowest MISE with single trials because it captures both transitions with only small attenuation. BARS emerges victorious when larger samples are used, but ARRIS performs reliably on even very small samples: In the single-trial cases, ARRIS sometimes makes attenuated estimates, but also sometimes does very well despite the paucity of data. As we will argue, it is this reliability in the face of small samples that makes ARRIS suitable for uncovering real-time changes in firing dynamics.

## Single Trial Rate Estimation

ARRIS and BARS are closely related, and both address the same problem: Fitting a model is hard when you do not know a *priori* how many parameters the model has. When the number and positions of a spline’s control knots are unconstrained, any arbitrary cloud of data can be fit exactly by an overfit curve with no subsequent predictive value. In order to make spline regression reliable as an inferential method, model selection procedures must severely constrain the number of knots in order to respect parsimony. Even with a handful of knots, however, discovering the optimal positions for those knots is an intractable problem, analogous to that of the traveling salesman.

BARS overcomes this difficulty by rendering the regression splines *adaptive*, using the reversible-jump Markov chain Monte Carlo algorithm (or “RJMCMC”) (Green, 1995). Like other MCMC algorithms (Geyer, 2011), RJMCMC uses stochastic methods to discover the contours of complex probability distributions. However, in addition to assessing the validity of various model parameter values, RJMCMC also dynamically adjusts the *number* of parameters, adding or removing them as part of its simulations (see also Fan, 2011). As applied to spline regression, RJMCMC will add, remove, and relocate knots dynamically, doing so in a manner that converges on the most parsimonious estimate. Once a stable estimate of the model is achieved, continued simulation maps out the sampling distribution of possible model estimates.

This technique is very effective in large datasets because the enormous flexibility displayed by splines is counterbalanced by RJMCMC’s natural tendency to discard needlessly complex models. However, when limited evidence is available, BARS favors highly attenuated models. This is demonstrated by Figures 1 and 2: Although abrupt transitions are evident upon visual inspection, BARS favors gradual transitions. Consequently, a different model specification paradigm is required to obtain reasonable estimates for small samples.

The simplest alternative to using splines is to employ a model consisting only of knots, with no smoothing whatsoever. In the models that ARRIS considers, “knots” cease being control points and instead become the coordinates at which an abrupt discontinuity occurs. Thus, activity is modeled using *only* discontinuous changes in rate. The sampling distribution of models is estimated using RJMCMC, just as it is for BARS. Importantly, however, the aim of ARRIS is not to find the single best series of step functions, but instead to average over the full sampling distribution of step functions produced by RJMCMC. Averaging over many different potential knot locations permits ARRIS to identify gradual changes in rate. This provides an advantage in small samples: The model average produced by ARRIS has the ability to fit both gradual and abrupt transitions even when data is scarce.

Let *D*_*r*_ correspond to a time series of events occurring during trial *r*. Each individual time bin *d*_*t,r*_ consists of a count of the number of spikes observed during that interval (here assumed to be a discrete 10ms block); typically, *d*_*t,r*_ will equal 0, 1, or occasionally 2. For the purposes of this analysis, we will assume that spikes arose from a Poisson process with a firing rate *λ*_*t,r*_ at time *t* on trial *r*. Although this assumption can be problematic, such problems typically appear in the spectral statistics of very short time scales (Lindner, 2006). The ARRIS procedure uses RJMCMC to adaptively sample the various possible step-wise models. For every coordinate in the dataset, this yields a sampling distribution *S*_*t,r*,1_. The mean of this sampling distribution is denoted by *μ*_*t,r*,1_. Thus:

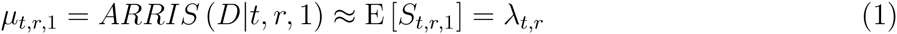

Here, the subscript 1 indicates an estimate based on a single trial.

Figure 3 provides an example of ARRIS performing a rate estimate given two separate trials of simulated data. A Poisson process generated 116 spikes (Figure 3A) and 90 spikes (Figure 3B) on two trials, according to the firing rate plotted as a black dashed line. These spikes are depicted as hash marks along the top of the figure. The best estimate of the firing rate, according to ARRIS, is plotted as a solid line. However, this best estimate is merely the mean of a sampling distribution at each point; also depicted are the 80% and the 99% credible intervals for the estimate, based on the corresponding upper and lower percentiles of *S*_*t,r*,1_.

**Figure 3:**
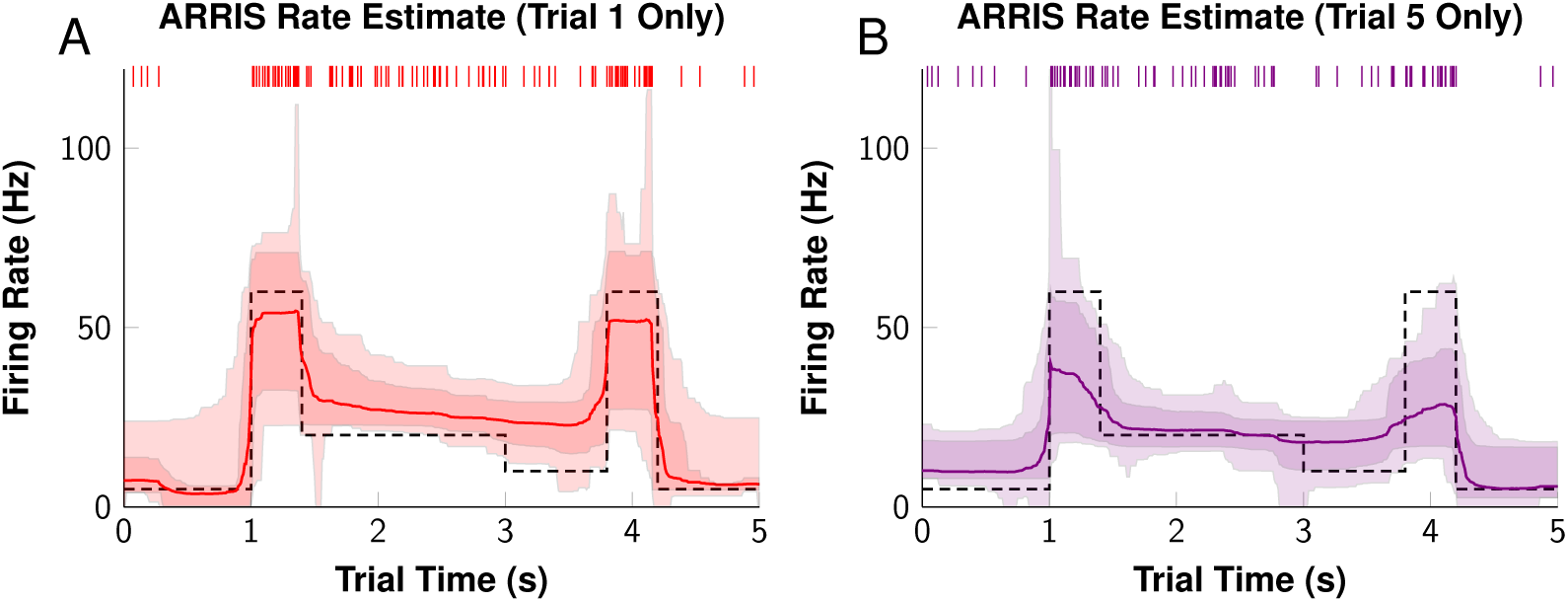
Single trial ARRIS estimates. Spike trains were generated according to the inhomogeneous Poisson firing rates (dashed lines). These spike trains are plotted along the top of each graph as hash marks. ARRIS was subsequently used to obtain estimates (solid lines) for the firing rates. In addition to the estimates, the 80% credible intervals (dark overlay) and 99% credible intervals (light overlay) are included to display estimate uncertainties. Because these estimates arise from single trials, they are very uncertain and, in some cases, far off the mark. In addition, subtle effects (such as the transition at 3s) cannot be detected based on a single trial.

The estimates in Figure 3 are noisy and cannot detect subtle changes (such as the shift in rate at 3.0s). Nevertheless, it performs reasonably well in identifying the major features of the firing rate, despite being based on a single trial in each case. In addition, the true firing rates are reliably within the credible intervals.

## Firing Dynamics That Change Over Time

The standard approach in rate estimation is to presume that every trial is exchangeable with every other trial, and to interpolate an estimate across all trials (in the style of Figure 1). Any method that relies on this logic makes a strong assumption that firing rates are stable over the observed period, and that any features revealed in the session average are representative of individual trials.

Figure 4 presents two simulated examples of how this “assumption of static activity” can lead to invalid estimates. In the left case, activity rises and falls during each trial, with the peak gradually shifting over consecutive trials. The shifting peak can be seen clearly in estimates taken from subsets of trials, but disappears in the marginal estimate. The right simulation also has only one peak per trial, but the position of this peak shifts suddenly between trials 25 and 26. Although analyses performed on subsets of trials show these distinct peaks clearly, the marginal estimate suggests that there are two peaks. Thus, the unjustified assumption of stable activity during a session can both conceal patterns of activity, or can produce a session average that does not resemble any of the individual trials.

**Figure 4:**
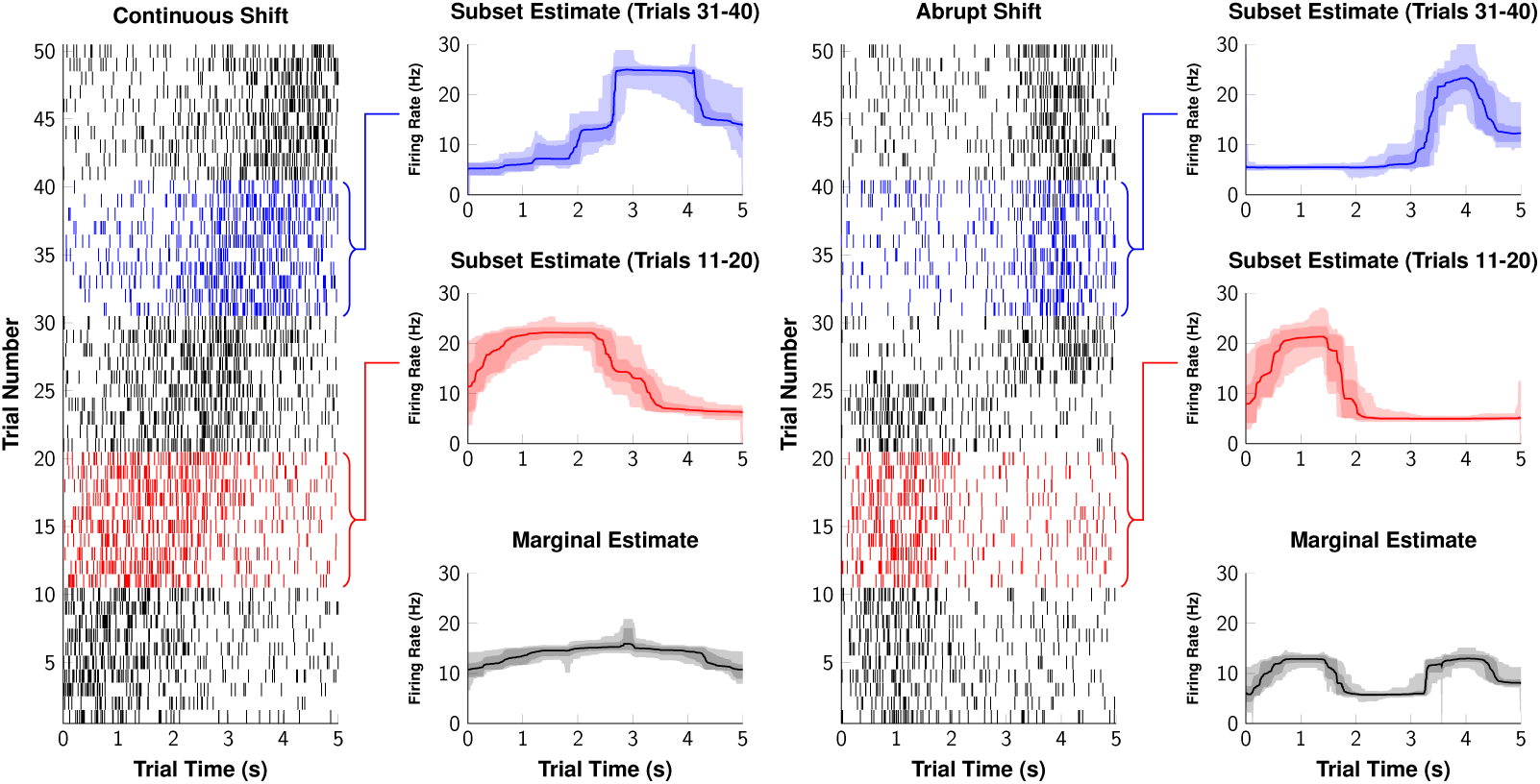
Firing rate estimation as firing dynamics change. Two simulated sessions of neural firing in which the pattern of firing changes over the course of the session. Shaded areas depict the 80% and 99% credible intervals for each estimate. **Left**. Although each trial has a peak of activity, the time of the peak drifts from early in the trial to late over the course of the session. Estimates performed on subsets of the trials (in red and blue) show these shifting peaks clearly. A marginal estimate using the entire session, however, shows no change in the firing rate, effectively masking this effect. **Right**. Again, each trial has a single peak of elevated activity. This peak occurs early in the trial for the first half of the session, then abruptly changes to appearing late in the trial. Subsets of trials (red and blue) can correctly characterize the single peak observed during the epoch from which they are sampled, but the marginal estimate predicts two peaks.

The examples provided in Figure 4 show clear changes over time on vi-sual inspection, but visual inspection alone does not provide a solution to the problem of which trials the analyst should group together. Furthermore, important differences may nevertheless be too subtle to spot visually. Admitting the need to analyze subsets of trials is not enough; a systematic approach to the problem is required to ensure that decisions of how to subdivide the analysis are driven by the evidence, and not by subjective interpretation of a visual impression.

When undertaking an analysis of subsets of trials, the primary tension that must be resolved is between undersmoothing and oversmoothing. Performing the analysis on very small sets of trials ensures that local features in the activity can be discovered, but each estimate will be very uncertain because each subset provides so little evidence. On the other hand, working with large sets of trials (e.g. first half of the session vs. last half) risks producing erroneous marginal estimates for the same reasons as full-session averaging.

One solution is to perform estimates using a moving window (e.g. an estimate using trials 1-11, then another using trials 2-12, etc.). However, selecting the width of the window is itself an optimization problem. If an overly wide window is chosen, then the estimates will be badly attenuated (as in Figure 1, row 1); if an overly narrow window is chosen, estimates will be noisy (as in Figure 2, row 1). Additionally, it is reasonable to assume that if trials 2 and 12 relate to an estimate centered at 7, then trials 1 and 13 probably also have some bearing on the estimate. At the same time, if an estimate is centered on trial 6, trials 5 and 7 are probably more similar than are trials 2 and 12. In other words, when interpolating an estimate over multiple trials, it is reasonable to assume that trials closer to the estimated time point are more relevant to the estimate than trials that occurred much earlier or much later in the session. This motivates the use of *weighted ensembles* of trials, with trials nearest to the time of interest contributing more evidence than those that are not as proximate.

Figure 5 presents a simulated example of firing that changes over time. Spikes are reported over a series of 50 trials, but the activity regularly changes its character. During the first 10 trials, activity is limited to two narrow intervals. Then, from trial 11 to trial 40, a broader unimodal distribution of spikes is evident, which gradually increases its spread over the interval, then contracts again. Finally, trials 41 to 50 return to restricting activity to two intervals.

**Figure 5:**
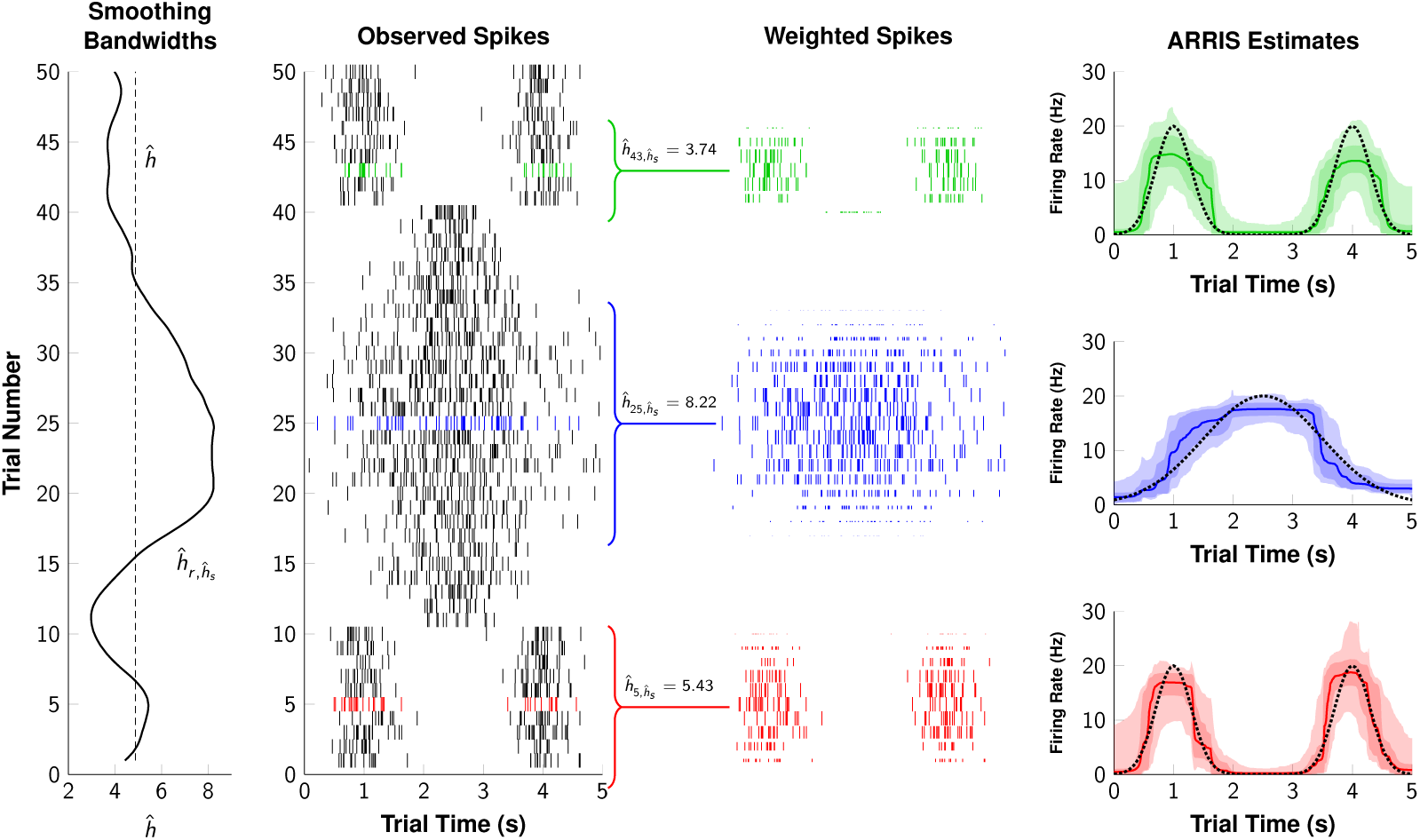
Local estimates of firing rate using variable smoothing bandwidths. **Left**. Optimal fixed (dashed line) and variable (solid line) bandwidths, as determined by optimizing GCV. **Left Middle**. Spikes observed during 50 trials of a simulated data set. **Right Middle**. Spikes near each of three demonstration trials (trials 5, 25, and 43). The height of the hash mark corresponds to the weight each spike contributes to the estimate (with very small spikes making very small contributions). **Right**. ARRIS estimates of activity for each demonstration trial (colored lines), along with 80% and 99% confidence intervals (shaded areas) and the original rate parameter used to generate the data (black dashed lines). Even when the bandwidth is so narrow as to include only a handful of trials, the original patterns of activity can be reliably recovered.

Given firing dynamics like those in Figure 5, it would be impossible to correctly describe the activity by simply marginalizing over all trials. However, a fixed-width moving window (e.g. an analysis of 10 trials at a time) will always blend dissimilar portions of the data (e.g. the discontinuity near trials 10 and 40). Using weighted interpolation, activity may be sampled from trials within some ‘bandwidth.’ For example, the estimate at trial 25 (in blue) uses a bandwidth of 8.22. Although fifteen different trials make a contribution to the estimate, their total weight is only that of about 9.5 trials (because the trials further from trial 25 count for much less than their full value). However, the estimate at trial 5 (in red) uses a bandwidth of 5.43, because a wider bandwidth would begin to include contributions from the unimodal middle epoch. This yields an estimate based on what are effectively 6.3 trials of evidence. When very close to a boundary (as with trial 43, in green), even narrower bandwidths are needed to avoid including spikes from a dissimilar epoch.

In order to obtain reliable estimates of firing in highly dynamic scenarios such as this one, several ingredients are needed:

- A weight function is needed to quantify how “close” any given trial is to a target time. We have chosen to use the tricube weight function (Cleveland, 1979).
- An bandwidth parameter *h* that governs the width of the weight function. Although an ‘optimal’ value 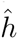 may be identified that does the best job across the session, Figure 5 demonstrates that no single bandwidth will be ideal for the entire session.
- Thus, each trial, in effect, needs its own bandwidth. Borrowing a strategy from Shimazaki & Shinomoto (2010), we specify a second optimized ‘smoothing bandwidth,’ 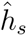, which ensures that bandwidths change smoothly over time.
- In order to evaluate the optimality of any given bandwidth *h*, we need a model selection criterion that maximizes the quality of our out-of-sample prediction. We use the generalized cross-validation index (GCV) (Golub et al., 1979), selecting values of *h* that minimized GCV.
- Iteratively minimizing GCV is computationally expensive. To facilitate bandwidth identification, a kernel density approximation to ARRIS is specified. This permits the discovery of relatively similar epochs of activity to be identified without requiring repeated ARRIS reanalysis for every candidate combination of bandwidths.

## The Tricube Weight Function

Interpolation over time was accomplished by giving each trial a weight using the tricube weight function (Cleveland, 1979):

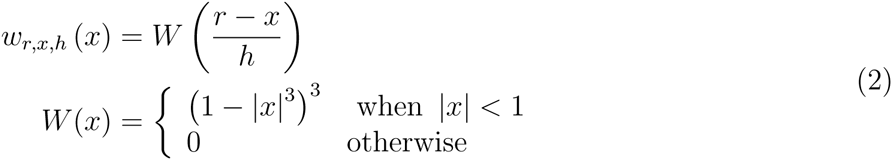

Here, *h* corresponds to the bandwidth of the weight function, *r* corresponds to the current trial that is being estimated, and *x* corresponds to some other trial whose weight needs to be computed. Every observation *d*_*t,x*_ makes no contribution to the local estimate when |*r-x*| > *h*; the rest make a weighted contribution, based on their proximity to *r*.

From these, a weighted mean of the firing rate may be computed from the data, denoted by 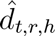:

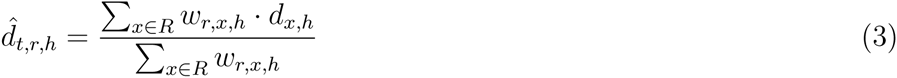

Rather than simply using *d*_*t,r*_ as the estimate of firing at time *t* on trial *r* (which is likely to be noisy), the estimate is instead the weighted mean of those trials closest to *r*, with only those trials within ±*h* making a contribution. We denote the estimate of the underlying rate given a bandwidth of *h* as *μ*_*t,r,h*_:

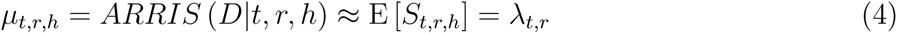

Note that, as defined, 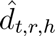 is not limited to integer values of *r*. If provided with a value of *r* that lies between any available observations, a weighted average will be computed in which none of the data receive a full weight of 1.0. This permits interpolation to arbitrary precision. As *h* → 1, the data used in the estimate converges on *d*_*t,r*_; As *h* → ∞, the data used in the estimate converges on the exchangeability assumption in which every trial in the session is included in the estimate and all are given equal weight.

## Optimizing the Bandwidth *h*

When performing the ARRIS procedure over a relatively stable interval, better estimates of the firing rate can be obtained by using larger values of *h*. At the same time, when ARRIS estimates activity during sessions in which rapid shifts in activity are observed, unattenuated estimates can only be achieved by using a small values of *h*. As with the kernel methods of Shimazaki & Shinomoto (2010), the choice of an appropriate value for *h* is a non-trivial optimization problem. There is, in principle, some value of *h* that best trades off the noisiness of localized samples of data with the attenuation that occurs when bandwidths include large swaths of the session.

A well-established model comparison metric that can measure this tradeoff is the generalized cross-validation score (GCV) (Golub et al., 1979). Ideally, systematic cross-validation of all subsets of the data reveals the degree to which a model reliably makes out-of-sample predictions, but such methods require factorial runtime. GCV provides a robust estimate of how well a given model would fare under cross-validation testing (particularly how overfit or underfit the model is) without the impossibly demanding computational burden of performing every cross-validation. GCV’s relative ease of computation has long made it an effective method for optimizing parameters that otherwise resist direct analysis (Craven & Wahba, 1979; Jansen et al., 1997; Josse & Husson, 2012; Jansen, 2015)

In order to identify a parsimonious value for *h*, one need only compute the GCV scores associated with various values and select the value of *h* that minimizes the GCV. This balances the goodness of fit associated with a given value of *h* against the roughness of the estimate:

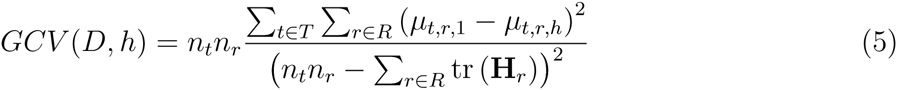

Here, tr (**H**_*r*_) corresponds to the trace of the hat matrix associated with the smoothing procedure on trial *r*, which provides an estimate of the effective number of parameters. *n*_*r*_ and *n*_*t*_ in turn correspond to the number of trials and the number of time bins, respectively. Because *GCV* is calculated across all *R* and all *T*, minimizing it yields a single value for *h*, applied across the entire spike raster. Henceforth let 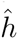 refer to the fixed bandwidth that minimizes GCV (and is therefore approximately optimal).

When the rate of change in the firing rate is steady from one trial to the next, 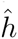 reasonably captures gradual changes. However, activity may change dramatically over a short interval, while also displaying long periods of stability in the intervening epochs. Consequently, it may be desirable to optimize *varying* values of *h*_*r*_ for each trial *r*. To do this, we adopt a strategy inspired by the approach to bandwidth optimization described by Shimazaki & Shinomoto (2010).

One approach that accomplishes this is to perform generalized cross-validation on each trial separately, optimizing a different *h*_*r*_ score for each by minimizing the corresponding *GCV*_*r*_ score:

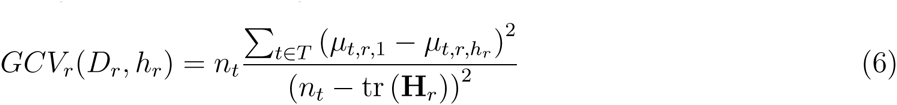

The value of *h*_*r*_ that minimizes *GCV*_*r*_ is considered optimal, and is denoted by 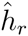. However, *GCV*_*r*_ depends on far fewer data than *GCV*. As a result, 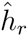 scores fluctuate considerably from one trial to the next as a function of sampling error. It is therefore desirable to specify an additional smoothing factor *h*_*s*_, which is used to smooth the volatile series of 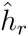 values as a function of their weighted geometric mean.

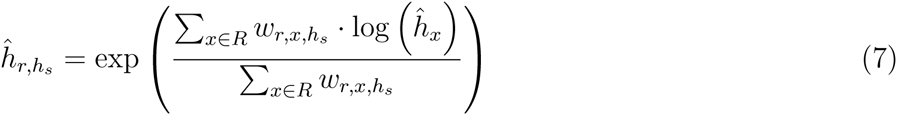

In order to obtain an optimal value for *h*_*s*_, generalized cross-validation may again be used:

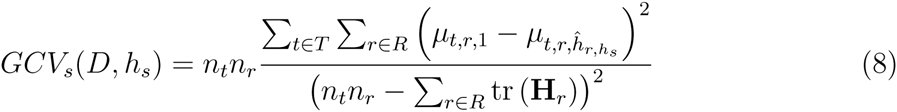

To summarize: In order to optimize a variable bandwidth, one must first minimize the *GCV*_*r*_ for each trial *r* in order to identify the optimal value of 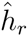. Then, this noisy vector of smoothing factors must itself be smoothed by using an optimized bandwidth 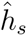. This, in turn, is identified by minimizing *GCV*_*s*_.

Because RJMCMC is a Monte Carlo method, its estimates are inevitably noisy and its approximated posterior distributions are jagged. This is true even when they are expected to be asymptotically smooth. As a final smoothing procedure, the resulting estimate should be smoothed with a bandwidth of 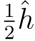 (for fixed bandwidths) or 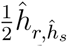 (for variable bandwidths). Because this final smoothing is done at a much finer grain than the original estimation procedure, it results in only minimal attenuation of the main signal.

In addition to serving a practical function, the time series 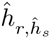 can also be considered a non-parametric measure of the speed with which patterns of neural activity change over time. Low values of 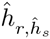 are needed when firing is changing rapidly. Examining how 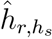 changes over time should thus reveal when adaptation occurs, as well as its relative speed.

## Rapid Evaluation of *h*

Unfortunately, optimizing *GCV, GCV*_*r*_, or *GCV*_*s*_ directly is extremely inefficient. Estimates of *μ*_*t,r,h*_ depend on RJMCMC, and consequently fluctuate slightly from one simulation to the next. This precludes the use of greedy hill-climbing algorithms. Although more error-resistant methods, such as simulated annealing, can overcome this problem, RJMCMC remains computationally intensive. Consequently, it is desirable to identify an alternate method for evaluating different values of *h* that yields reasonable results, but does so rapidly and reproducibly.

One such approximation is offered by kernel estimation procedures (Bowman & Azzalini, 1997). Given a particular kernel bandwidth, the resulting estimate can be rapidly computed. Furthermore, since kernel methods do not rely on Monte Carlo methods, they yield identical results every time they are re-run, enabling hill-climbing algorithms to converge on optimal GCV scores without being mislead by local minima.

In order to best approximate ARRIS estimates, within-trial firing rates were estimated using a box kernel with a bandwidth of 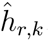. Rather than optimize this kernel bandwidth, we used a plug-in estimator. The most popular such estimator is Silverman’s rule-of-thumb (Silverman, 1986), which is defined in terms of the sample size and the estimated standard deviation:

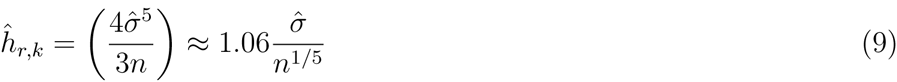

We have presumed that the spikes are generated by something approximating a Poisson process with a rate parameter λ. Poisson processes are expected to have inter-spike intervals that follow an exponential distribution whose parameters are also governed by λ. Using this relationship, we anticipate how many spikes we should expect during an interval of time (effectively, an estimate 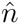), and how dispersed those spikes should be (an estimate 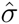). Both estimates then can be combined to provide our plug-in estimator of the bandwidth.

Let Δ*D*_*r*_ denote the set of intervals between consecutive spikes during trial *r*. Under the assumption that these inter-spike intervals are exponentially distributed, a robust estimator of the rate is provided by 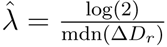, where mdn() is the median (Forbes et al., 2011). Using the median, rather than the mean, prevents the estimate from being overly influenced by long periods of inactivity that might occur between clusters of spikes. Having an estimate of 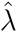 then provides an estimate of the standard deviation 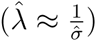 and of the expected sample size 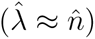.

Since 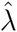 governs both the dispersion of spikes and their expected frequency over each unit of time, we can substitute it into Silverman’s rule of thumb to yield the following plug-in estimator of the bandwidth:

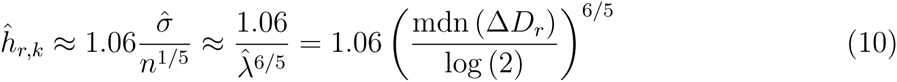

Let *ĸ*_*t,r,h*_ correspond to a kernel estimate of the firing rate, using a box kernel with the above bandwidths 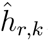. This estimate is analogous to the value *μ*_*t,r,h*_ obtained using the ARRIS procedure. When *h* = 1, this corresponds to ordinary kernel density estimation. When *h* > 1, the observed spikes differ in their weights, requiring a weighted kernel density estimate to be computed (Guillamón et al., 1998). In either case, this permits rapid approximation of the corresponding *GCV* scores:

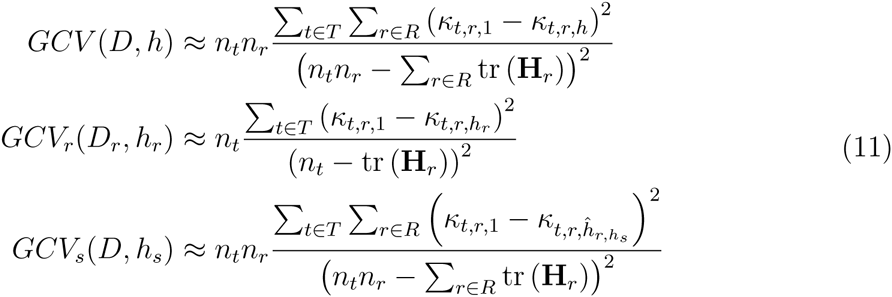

When values of *h, h*_*r*_, and *h*_*s*_ are selected based on optimizing box kernel estimates rather than ARRIS estimates, they respond to the same general features of the data that ARRIS would respond to. When the session is generally stable (i.e. when consecutive trials resemble one another), higher values of *h* will be favored. When there are dramatic changes, lower values of *h* will instead be favored to avoid mixing trials that are dissimilar. Once the kernel approximation has provided a map of how narrow or broad the weighted interpolation needs to be, ARRIS then goes on to provide a more reliable estimate using the smoothed values of 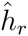.

The principle application of the ARRIS procedure is to characterize changes in the firing rate over the course of a recording session. In order to demonstrate the efficacy of our approach, we first present two examples using simulated data. This permits comparison of the estimates to the true underlying rates.

In our first simulation (introduced in Figure 5), spikes were generated by a Poisson process whose rate *ρ* was governed by the following function, measured in Hz:

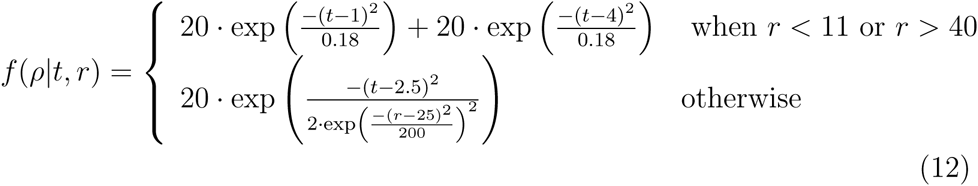

Thus, during the first 10 trials and the last 10 trials, firing was governed by two static Gaussian functions. During the intermediate period, firing was governed by a single Gaussian function whose standard deviation varied as a function of trial number. A complete map of the estimated firing rate is plotted in Figure 6A, and the resulting spike raster is plotted in Figure 6B. Here it becomes especially clear that the variable bandwidths capture the broad strokes of the firing reliably, despite the rapidly changing character of the underlying generating function.

**Figure 6:**
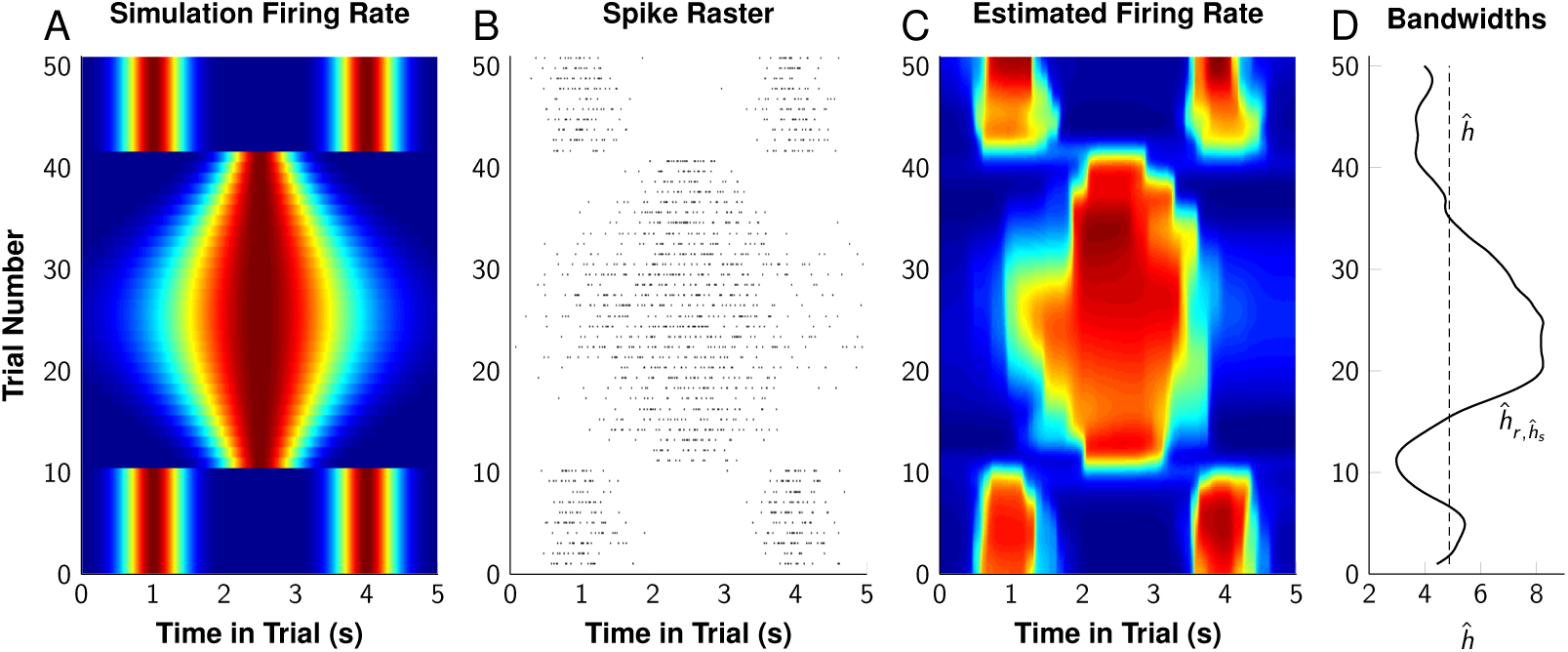
Simulation of discontinuous multimodal firing. 50 trials of simulated spike trains were generated using a function that changed over time. **A**. The veridical firing rate, varying from 0 to 20 Hz, over the 50 simulated trials. **B**. The spike raster resulting from the simulation. **C**. The ARRIS estimate of the firing rate, using optimized dynamic bandwidths 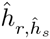. **D**. The optimal fixed (dashed line) and dynamic (solid line) bandwidths for these data, as determined using kernel density approximation and generalized cross-validation.

As previously described, Figure 5 depicts how estimates were obtained for three of these simulated trials. The optimized variable bandwidth favored a wider sample of trials near trials 25 because the data appeared more stable in that interval, whereas much narrower bandwidths were favored near transitions. The tricube weight function further ensured that each local estimate was primarily influenced by those trials near the trial of interest, with distant trials making only small contributions to the estimate. These permitted, within a single analysis, the detection of the three different epochs of firing.

Optimal bandwidths were obtained for this dataset using kernel density approximation. This yielded an optimal fixed bandwidth of 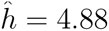 and an optimal smoothing bandwidth of 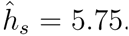. Figure 6C shows both the estimated firing rate, while Figure 6D shows the varying values of 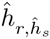. Despite being presented with data that (1) display multiple abrupt transitions in the underlying generating function and (2) show dynamic change in the generating function, ARRIS nevertheless captures the major patterns of activity successfully. A factor contributing to this recovery is the large variation in the values of 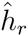 over the course of the session. By displaying narrow bandwidths near the discontinuities after trials 10 and 40, but broad bandwidths during the comparatively stable firing near trial 25, ARRIS is able to approximate both varieties of features.

To examine this estimate in more detail, Figure 7A-C plots the estimates during three different trials: 5, 25, and 45. The estimated function (black line) compares very favorably with the generating function (dotted line) in every case. Additionally the 80% credible intervals (dark gray) and 99% credible intervals (light gray) are shown for each estimate, and the true rate reliably falls within those bounds during all but a handful of time bins.

**Figure 7:**
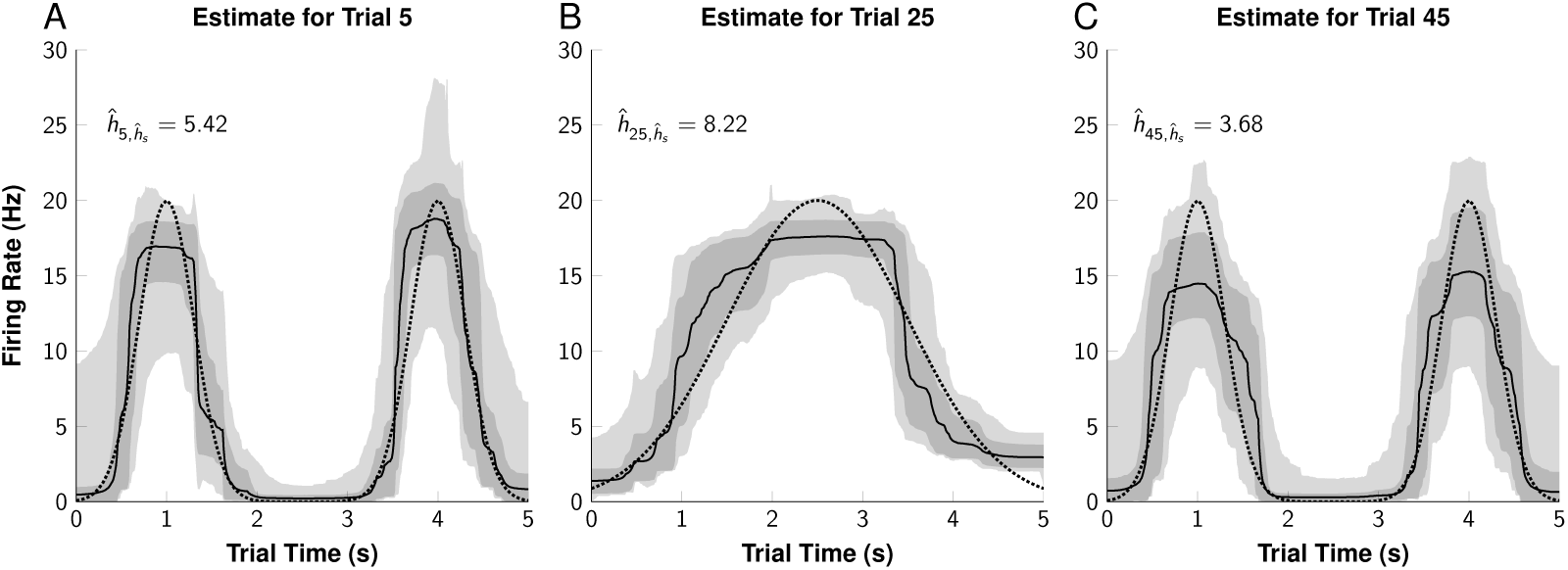
ARRIS estimates of individual simulated trials. Estimates of the firing rate during three trials (5, 25, 45), given an optimized dynamic smoothing bandwidth. The firing rate responsible for the data is shown as a dotted line, whereas the ARRIS estimate is shown as a solid line. The dark and light shaded overlays represent the 80% and 99% confidence intervals, respectively.

Although the credible intervals consistently include the true rate, the estimate is somewhat attenuated near the peaks of the generating function. This can be credited in part to the small amount of data in this example. Given that this simulated example makes use of only 50 trials, and that the frequent discontinuities in firing require a very narrow bandwidth, each of the estimates in Figure 7 is based on the weighted contribution of less than ten trials. Under these circumstances, the information available from the spikes is insufficient to capture those fine details. Consequently, it is important to report not only the estimate, but the credible interval established from the sampling distribution *S*_*t,r*_.

Another manner in which the estimates may be evaluated is across trials during a given window of time. Figure 8A-D does so at four trial times: 1s, 2s, 3s, and 4s. The estimate reliably hews close to the true rate, keeping it within the 80% credible intervals (dark gray) and 99% credible intervals (light gray). Even the sharp discontinuities at trials 10 and 40 yield fairly rapid adjustments, although these transitions do display slight attenuation.

**Figure 8:**
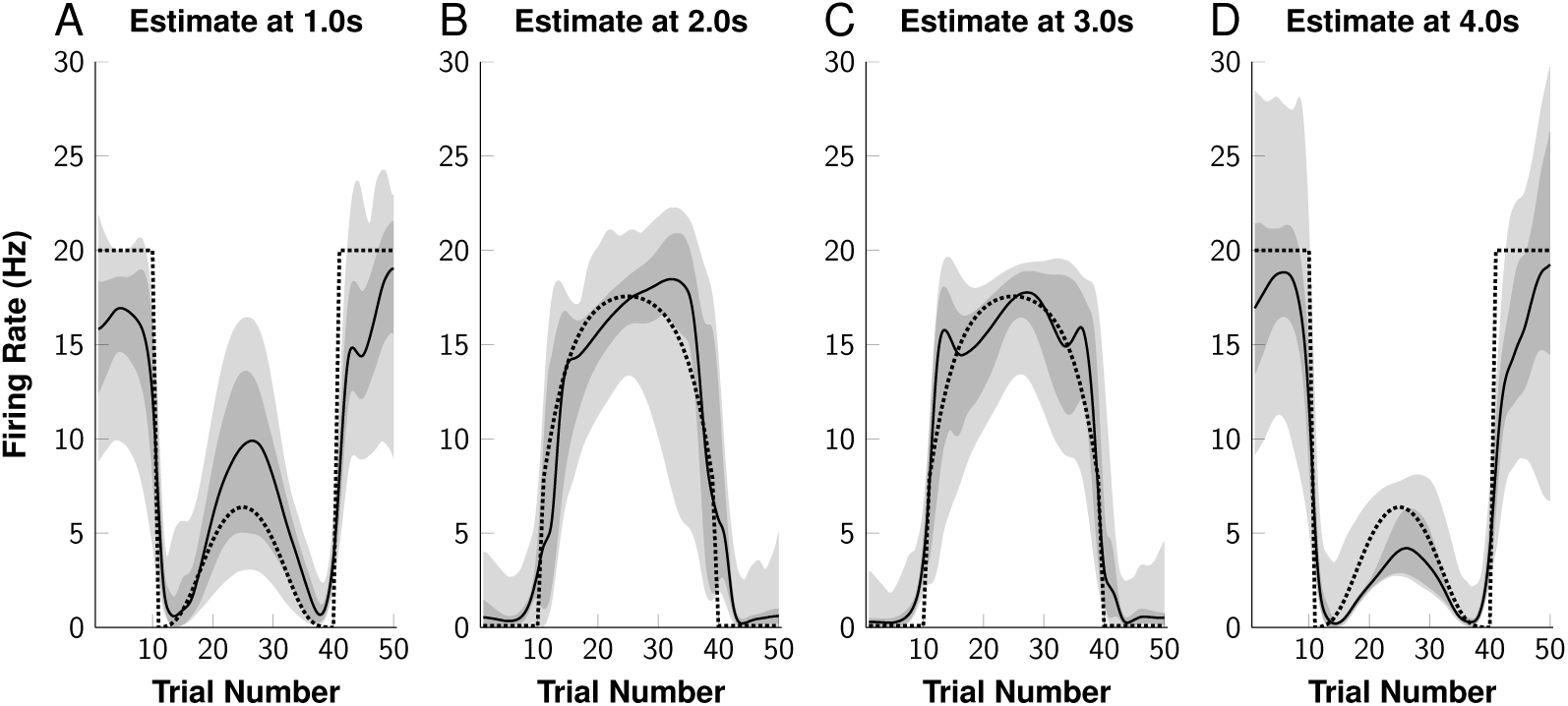
ARRIS estimates at specific times across trials. Estimates of the firing rate at four different trial times (1s, 2s, 3s, 4s) across the full session, given an optimized dynamic smoothing bandwidth. The firing rate responsible for the data is shown as a dotted line, whereas the ARRIS estimate is shown as a solid line. The dark and light shaded overlays represent the 80% and 99% confidence intervals, respectively.

The estimates presented in Figure 8 are smooth even between trials. For example, estimates were obtained not just for trials 1 and 2, but for the intervening values of 1.2, 1.4, 1.6, and 1.8 as well. The ARRIS procedure can be used to make estimates for arbitrary values of *r*, even those in between integer values.

Our second simulation demonstrates a more ambitious objective: To reconstitute the underlying changes in firing given only a subset of the available data. Figure 9A presents the source function of a Poisson process, with rate *ρ* (in Hz), governed by the following equation:

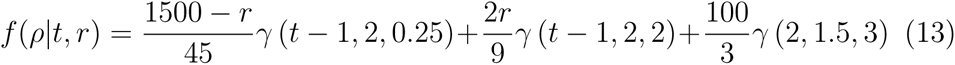

**Figure 9:**
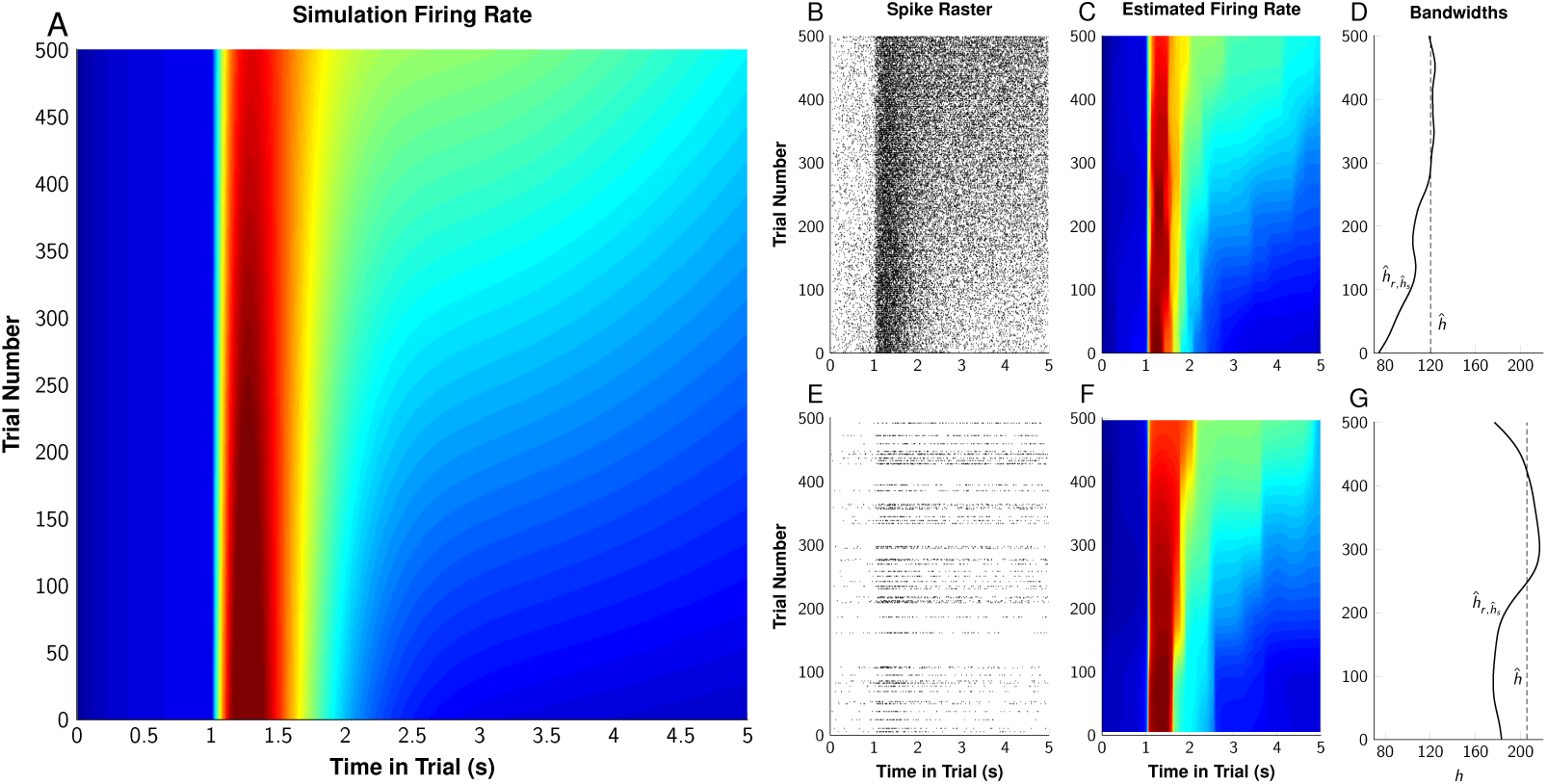
Simulation of smoothly transitioning firing. 500 trials of simulated spike trains were generated using a function that changed over time. **A**. The veridical firing rate, varying from 0 to 55 Hz, over the 500 simulated trials. **B**. The spike raster resulting from the simulation. **C**. The ARRIS estimate of the firing rate, using optimized dynamic bandwidths 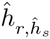. D. The optimal fixed (dashed line) and dynamic (solid line) bandwidths for these data, as determined using kernel density approximation and generalized cross-validation. E-G. Spike raster and ARRIS analysis of a randomly selected subset of 10% of the original data (50 trials).

This function was used to generate a spike raster for 500 trials (with trial number denoted by *r*) lasting 5 seconds (with trial time denoted by *t*). The full spike raster is presented in Figure 9B. From these 500 trials, we selected 50 of the trials at random; these spikes are plotted in Figure 9E. In addition to using the interpolation procedure to smooth out the error in estimation of the full dataset, we also attempted to reconstruct the firing using the 50-trial subset.

Figure 9C presents the interpolated estimate of the firing rate based on the full 500 trials, and Figure 9D shows the corresponding values of the optimized bandwidths. Figure 9F, in turn, presents the estimate based on the 50-trial subset, and Figure 9G shows the bandwidths optimized for that subset of data. Even with a minority of the trials, the overall form of the source function can reliably be recovered. In both analyses, the event-related potential after the 1s mark can be seen to decline, whereas the period following the peak gradually rises.

Note that the rate estimates provided by Figure 9F are continuous over the entire session, despite the large irregular gaps in the spike raster. For example, even though there was a wide gap between trials 110 and 160, weighted interpolation nevertheless permits an estimate for any one of those trials, and yields an estimate comparable to that obtained when all trials were used (albeit with additional uncertainty). Larger values of 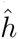 or 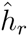 are generally required when working with a subset of the full session, as can be observed by comparing the optimized bandwidths in Figure 9D and Figure 9G.

Figure 10A-C shows the fit for three specific trials, based on the full data (top row) and on only 50 trials (bottom row), along with the 80% credible interval (dark gray) and the 99% credible interval (light gray). When all the data are available, the source function can be recovered almost exactly, differing only in the difficulty ARRIS displays capturing the apex of the peak. In the case using only 10% of the data (Figure 10D-F), the mean rate reasonably approximates the source function, but broad attenuation effects persist in the 99% credible intervals. In both cases, ARRIS succeeds at modeling the abrupt discontinuity at the 1s mark.

**Figure 10:**
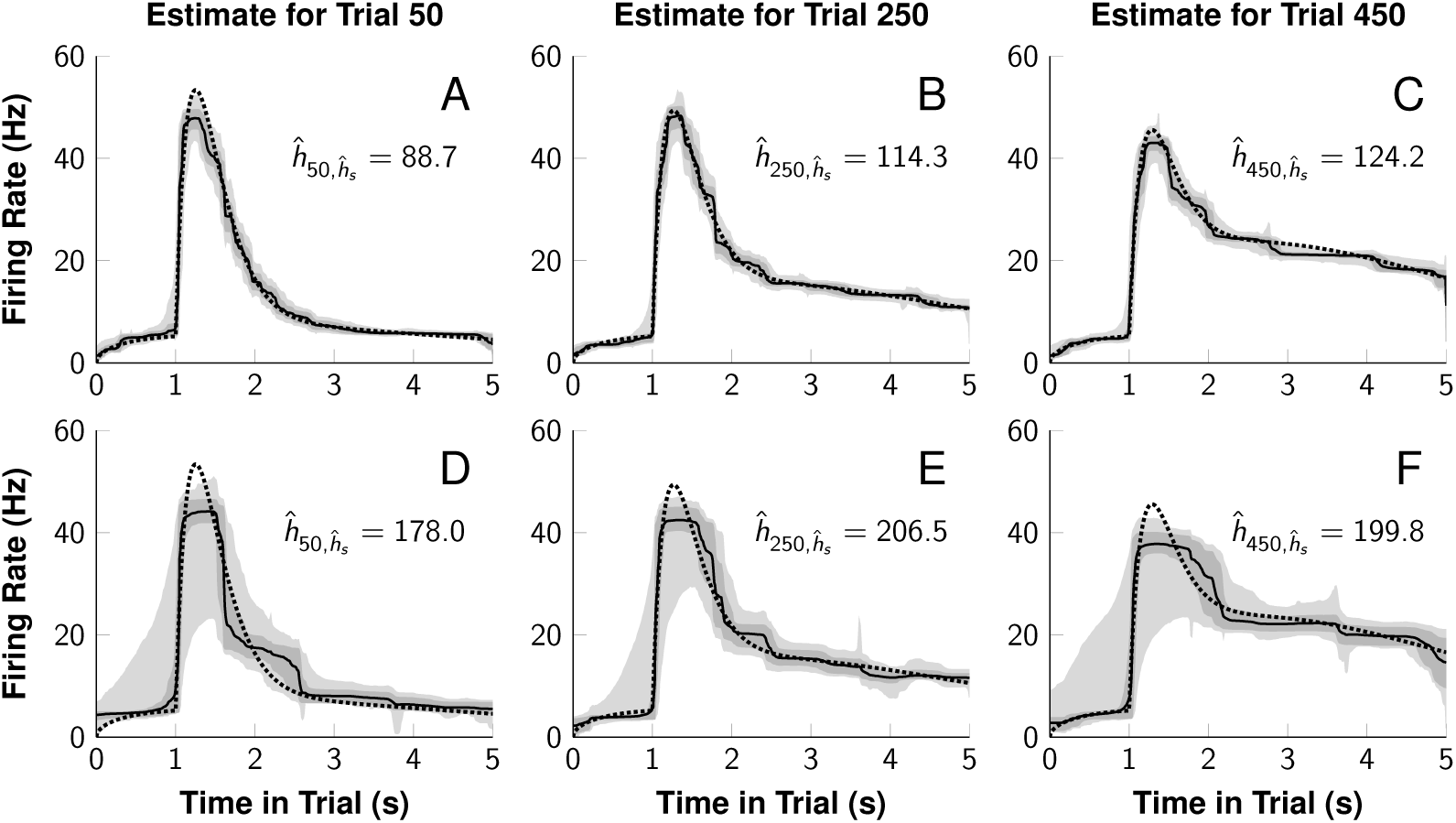
ARRIS estimates of individual simulated trials. **A-C**. Estimates of the firing rate during three trials (50, 250, 450), given an optimized dynamic smoothing bandwidth. The firing rate responsible for the data is shown as a dotted line, whereas the ARRIS estimate is shown as a solid line. The dark and light shaded overlays represent the 80% and 99% credible intervals, respectively. **D-F**. Estimates of the same trials, this time based on a random subset of 10% of the trials. Because ARRIS performs a principled interpolation over gaps in the data, these estimates may be obtained and their credible interval evaluated even when these trial numbers are not included in the 10% subset.

To demonstrate learning in an empirical case, ARRIS was used to estimate firing rates during a transitive inference procedure (Jensen et al., 2013). In the transitive inference paradigm, subjects are shown pairs of stimuli, drawn from an ordered list. Choosing the stimulus positioned earlier in the list results in a reward. Subjects are naive about the list ordering at the beginning of a session: Over successive trials, they must deduce the full ordering of list items on the basis of these pair-wise comparisons.

A rhesus macaque was presented with pairs of images draw from a 7-item list. Choices were made using eye movements, using equipment and procedures described by Teichert et al. (2014). Trial times were centered at stimulus onset (zebra stripe). Some trials were omitted because the subject did not fixate on the start stimulus (e.g. near trial 460). Despite these gaps, ARRIS had no difficulty producing an interpolated estimate of activity.

Figure 11 presents both a spike raster and an ARRIS estimate of activity in a parietal cell, showing a pronounced visual response that grew over successive trials. Although traditional analyses would reveal a strong visual response, their assumption of trial exchangeability would obscure this steady increase in the rate of firing.

**Figure 11:**
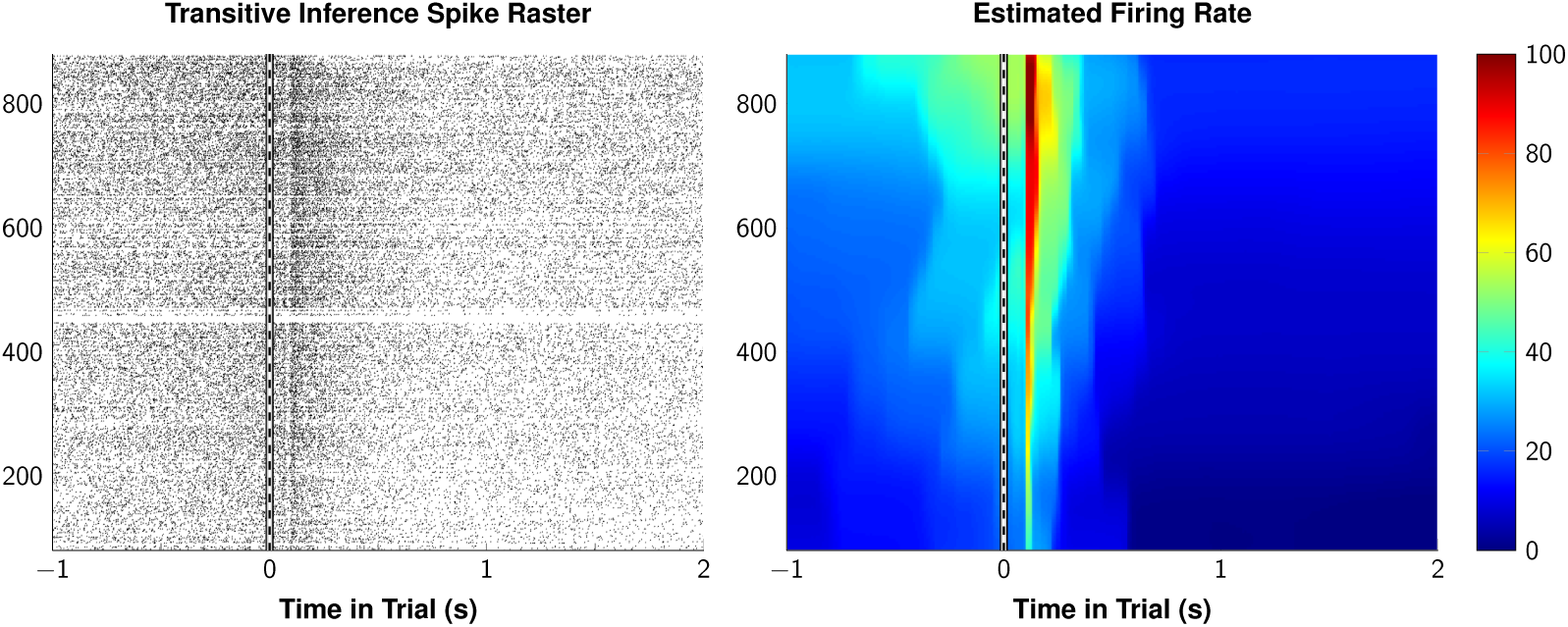
Estimated dynamics of firing across all trials of a transitive inference task. A rhesus macaque was presented with the 21 possible pairs of stimuli from a 7-item list. The subject was unfamiliar with the stimuli at the outset, and learned the ordered list by trial and error over the course of the session. **Left**. Raster of spikes from a parietal neuron during completed trials. **Right**. ARRIS estimate of firing, interpolated across all trials.

Changes in firing that emerge during this process can be compared with performance to provide insight into the dynamics of learning. Figure 12 shows the correspondence between peak neural activity (occurring approxi-mately 120ms after stimulus onset) and overall response accuracy. Even at the beginning of the session, the stimulus onset evokes a clear 60 Hz response from the cell. However, as accuracy gradually rises to approximately 85%, the firing rate rises to as high as 110 Hz.

**Figure 12:**
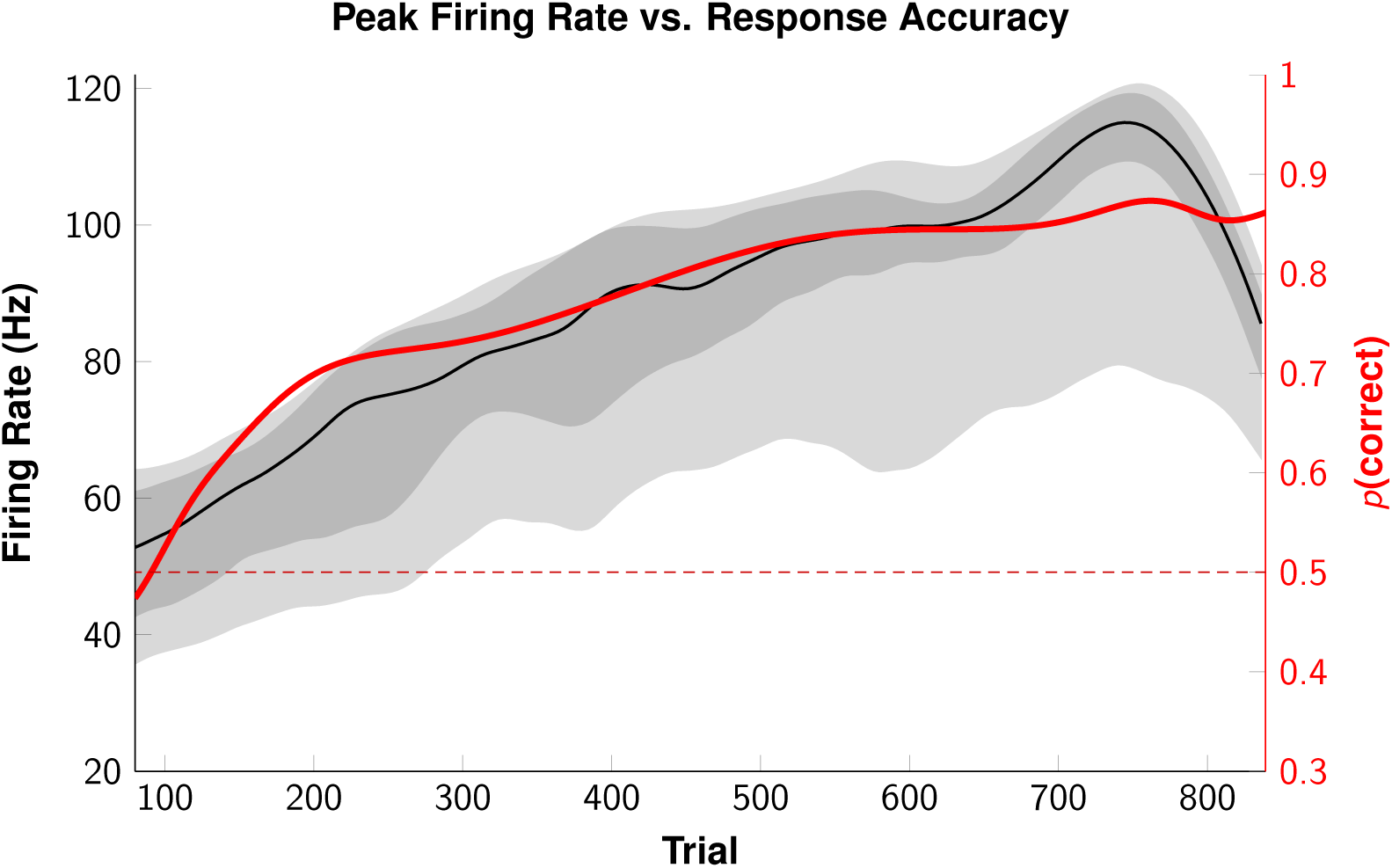
Estimated dynamics of firing as a function of response accuracy. The peak estimated firing rate from Figure 11 (black), compared to a smoothed estimate of response accuracy over the course of the session (red). Shaded regions correspond to the 80% and 99% confidence intervals, respectively. The dashed line corresponds to chance levels of accuracy.

## Discussion

ARRIS permits the firing rate of a neuron to be rigorously evaluated, even in small samples, thanks to the RJMCMC algorithm. This in turn provides a way to evaluate how activity changes over the course of a recording session. It provides an instantaneous estimate of any point during training by drawing evidence from a weighted ensemble of trials centered on the time of interest. The set of trials included, and their relative weight, is optimized to ensure that only small sets of trials are included when firing is changing rapidly, while larger sets of trials are used when activity is comparatively stable.

The major potential for ARRIS is to assess firing rates in naïve subjects. Historically, the norm in electrophysiology is to overtrain subjects in order to ensure that performance is at a stable ceiling. This, in turn, justifies the use of estimation procedures that require full exchangeability among trials. Because ARRIS can provide an optimized estimate of changing firing rates, even when those changes are relatively abrupt, fruitful electrophysiology can now be done *during* training. This will enable important questions to be asked, such as whether changes in performance can be detected before changes in activity.

The smoothed bandwidth used in weighted interpolation (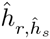) also provides a useful summary of changing activity, as it can be used as an indicator of how rapidly patterns of firing are changing. For example, in Figure 5, lower values of 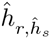 signal moments of abrupt transition, whereas higher values correspond to relatively stable firing. The slope of the rise and fall of 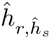 effectively models the acceleration and deceleration of change in the firing dynamics.

ARRIS is also useful when making estimates based on small samples, even when weighted interpolation is not strictly required. For example, Morrison et al. (2011) made use of a design in which subjects learned which stimuli were paired with rewards and punishments, only to have those contingent relationships suddenly reversed. Changes in firing rates caused by surprisal are likely to be brief, but could be characterized using an ARRIS analysis that was limited the trials immediately following each reversal.

The use of weighted interpolation not only presumes similarity between temporally proximate trials, but emphasizes similarity while also treating deviations as resulting from measurement error. This is necessary in neurons that are involved in learning and memory, but may not be appropriate in strictly sensory neurons that respond exclusively (and without adaptation over time) to properties of the stimulus. By treating session time as an important covariate, weighted interpolation can be used to rule out learning effects for neurons that do not display them. Having ruled out learning, ARRIS without weighted interpolation (as implemented in Figure 1) could then be used to obtain a reliable marginal estimate over the full session, as could more traditional methods of characterizing firing rate.

ARRIS is used for temporal interpolation, but it is intrinsically a single unit estimation procedure. Consequently, one open question is how best to apply ARRIS to simultaneous multi-unit recording. One possibility is to use weighted mixtures of multiple units to obtain population estimates. Thus, rather than mixing trials that were recorded at different times (as in Figures 11 and 12), a weighted mixture of multiple simultaneous units could be used. Such an analysis would, however, require solving the difficult problem of discovering the optimal weights by which to combine units. Thus, another more feasible approach would be to perform the ARRIS analysis on the individual recording units, then to use integrative techniques, such as representational similarity analysis (Kriegeskorte et al., 2008), to discover the patterns of firing among ensembles of neurons.

A limitation of the present implementation of ARRIS is that it does not incorporate refractory periods into the analysis. This is not a problem when smoothing over many trials, as the mixture of trials rapidly comes to resemble that of a Poisson process with respect to their inter-spike interval distributions (Hanes et al., 1995; Kass et al., 2005; Lindner, 2006). The concern remains, however, for single-trial estimates. Several potent models for refractory firing have been proposed, such as the inhomogenous Markov interval model (Kass & Ventura, 2001) and the point-process likelihood model (Truccolo et al., 2005). Both of these methods, however, are computationally intensive and data-hungry.

This raises the as-yet unresolved problem of how to best characterize firing patterns that violate Poisson assumptions, given very small samples. When spikes are assumed to be independent random events, the Poisson distribution is unambiguously the maximum entropy distribution that describes their frequency (Harremëes, 2001). However, when this assumption is relaxed, there are an infinite number of hypotheses explaining how spikes might be interdependent. Differentiating between such models is only possible with relatively large bodies of data. In light of the difficulty in distinguishing between models, the best general approach given small samples is to simply rely on descriptive inference using the negative binomial distribution (Pillow & Scott, 2012).

In conclusion, ARRIS has the ability to make sensible estimates of a cell’s firing rate given only a few trials. This, when combined with a strategy of weighted interpolation over session time or over perceptual space, makes it possible to obtain instantaneous estimates of activity. This will permit, for the first time, a systematic examination of how firing changes in response to real-time feedback from the environment.

## Code

Functions implementing the ARRIS algorithm are available on GitHub in the /belarius/matlab-ARRIS repository.

## Acknowledgments

The authors would like to thank Yelda Alkan, Benjamin Kennedy, and Orly Morgan for feedback during the development of this procedure, and for their assistance in data collection. This work was supported by US National Institute of Mental Health, grant number 5R01MH081153, awarded to Vincent Ferrera and Herbert Terrace.

